# Dynamics of brain activity reveal a unitary recognition signal

**DOI:** 10.1101/165225

**Authors:** Christoph T. Weidemann, Michael J. Kahana

## Abstract

Dual-process models of recognition memory typically assume that independent familiarity and recollection signals with distinct temporal profiles can each lead to recognition (enabling two routes to recognition), whereas single-process models posit a unitary “memory strength” signal. Using multivariate classifiers trained on spectral EEG features, we quantified neural evidence for recognition decisions as a function of time. Classifiers trained on a small portion of the decision period performed similarly to those also incorporating information from previous time points indicating that neural activity reflects an integrated evidence signal. We propose a single-route account of recognition memory that is compatible with contributions from familiarity and recollection signals, but relies on a unitary evidence signal that integrates all available evidence.

---

A repeated exposure to people or objects sometimes evokes only a vague sense of familiarity; at others, it elicits vivid recollections of contextual details from previous encounters. This distinction is formalized in dual-process models of recognition memory that posit two independent types of evidence subserving recognition decisions (with recollection commonly, but not always, conceptualized as a threshold process; Diana, Reder, Arndt, & Park, 2006; Yonelinas, 2002; Yonelinas, Aly, Wang, & Koen, 2010; Malmberg, 2008). In apparent support of these models, neuroscientific studies of recognition memory have identified patterns of brain activity with distinct time courses thought to reflect an early familiarity signal (peaking around 400 ms after onset of a memory probe) and a later recollection signal (peaking around 600 ms after probe onset; Curran, 1999; Rugg & Curran, 2007).

Most dual-process models assume that familiarity and recollection signals can each separately lead to recognition (Reder et al., 2000). In some models, however, the memory system integrates evidence from different sources into a unitary evidence signal (Rotello, Macmillan, & Reeder, 2004; Wixted & Mickes, 2010). This results in a single route to recognition despite the contributions from different types of evidence. From this perspective, such models are conceptually similar to single-process models which assume only a single evidence source (Malmberg, 2008). One indication that two separate routes to recognition may not be neccesary to account for recognition performance is the fact that single-process models have been highly successful at accounting for intricate relationships between response time distributions, accuracy, and confidence ratings across a wide range of experimental manipulations (e.g., Ratcliff, 1978; Ratcliff & Starns, 2009; Wixted, 2007; Dunn, 2004, 2008; Cox & Shiffrin, 2012; Diller, Nobel, & Shiffrin, 2001; Starns, White, & Ratcliff, 2012; Starns & Ratcliff, 2014; Shiffrin & Steyvers, 1997). Additionally, recent work linking brain activity to parameters in a single-process model of recognition memory has cast doubt on the common interpretation of early and late electrophysiological signals indexing familiarity and recollection respectively (Ratcliff, Sederberg, Smith, & Childers, 2016, but see also Anderson, Zhang, Borst, & Walsh, 2016, and Borst & Anderson, 2015, for an alternative approach that preserves this interpretation—we will return to this issue in the discussion).

Because the single- vs. dual-process labels do not reliably differentiate between the number of routes to recognition, we will refer to models as single- or dual-route models to make this distinction explicit. Specifically, we label models that assume that different types of evidence can give rise to different kinds of recognition decisions (e.g., Yonelinas, 1994, 1997; Reder et al., 2000; Diana et al., 2006) as dual-route models. Single-route models are those that assume a single type of evidence source (e.g., Shiffrin & Steyvers, 1997) and those that assume that evidence from multiple sources/processes is integrated into a unitary evidence signal (e.g., Rotello et al., 2004; Wixted & Mickes, 2010). Within the framework of dual-route models, it makes sense to label individual recognition decisions with respect to the type of evidence (e.g., “familiarity” vs. “recollection”) that gave rise to them, whereas such a categorization of individual recognition memory decisions is not meaningful within the framework of single-route models, because information from all available sources contributes to recognition decisions. We propose that conflating the question about the number of recognition signals (i.e., the distinction between single- vs. dual-process models) with the question about the number of different routes to recognition may have contributed to the apparent disconnect between the evidence for separate familiarity and recollection signals and the success of single-process models.

Capitalizing on the presumed temporal separation of familiarity and recollection signals (Diana et al., 2006), we quantify the neural evidence distinguishing targets from lures in various partitions of the period leading up to the recognition decision. Specifically, we ask if combining neural evidence from multiple time bins during the recognition decision tells us more about whether an item has been studied than just the latest considered time bin by itself. If we are picking up on independent signals at different points in the recognition decision, then combining information from both should boost our ability to use neural activity to distinguish between old and new items. If, however, the neural signal corresponds to an integrated/unitary evidence signal, information from previous time points should not contain information that is not also present in the neural activity at later points.

Figure 1 illustrates our approach with the help of two toy models of evidence in recognition memory. Figure 1A shows activation for two sources of evidence containing information about the old/new status of an item as a function of time, and Figure 1B shows two alternative ways these sources could give rise to an evidence signal for the recognition decision (in this toy example we assume an “early” and a “late” source, analogous to the presumed dynamics of familiarity and recollection signals). The top panel of Figure 1B illustrates a dual-route model: the recognition decision is based exclusively on whichever source has accumulated more evidence at the time of response. Thus any information from the non-dominant evidence source is lost. Assuming sources with different temporal signatures, the evidence signal will initially be determined by activity from the early source, which sometimes will be exceeded by activity from the later source by the time the response is initiated. The bottom panel of Figure 1B illustrates a single route model: here the evidence for the recognition memory decision at any given time reflects the information accrued across all sources so far. Even when the relative contributions of the different sources change, no information is lost, because all relevant information contributes to the evidence signal.

**Figure 1.**
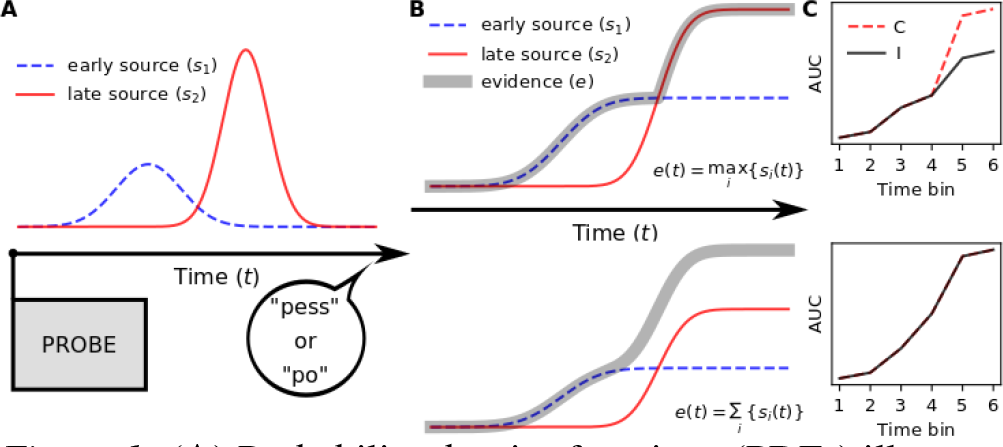
(**A**) Probability density functions (PDFs) illustrating two sources of evidence for recognition memory decisions. (**B**) Cumulative density functions (CDFs) for the PDFs shown in (**A**) along with CDFs for the evidence on which the recognition memory decision is based. The top panel illustrates a case where the evidence is determined by a single source of evidence leading to different routes to recognition memory depending on which source determines the evidence signal at the time of response (in the case of two sources, we label this class of models “dual-route models”). The bottom panel illustrates a case where the evidence signal integrates information from all sources (we label this class of models “single-route models” regardless of the number of sources contributing to the evidence). (**C**) Expected patterns of performance for classifiers trained on features from individual (**I**) or cumulative (**C**) time bins partitioning the time between probe onset and recognition response (see text for details). Assuming the sources contribute independent information with distinct dynamics, dual-route models predict diverging performance for classifiers trained on individual and cumulative time bins (top panel), whereas single-route models predict identical performance (bottom panel; C and I lines are overlapping). AUC indicates area under the receiver operating characteristic curve, a measure of classifier performance.

It is difficult to distinguish between these alternative accounts on the basis of recognition decisions alone, because these presumably only reflect a snapshot of the evidence signal from around the time when the response was initiated. Recordings of brain activity, however, allow us to assess the evolution of a neural evidence signal in the lead-up to a recognition response. We used multivariate (“machine learning”) classifiers to quantify the neural evidence distinguishing between targets and lures during the processing of the probe (i.e., between probe onset and just prior to the execution of a response). By comparing performance for classifiers trained on neural features from various partitions of this time period, we can make inferences about whether relevant information is integrated into a single evidence signal or whether evidence from an earlier signal is sometimes lost.

Figure 1C illustrates the logic of the main analyses. As explained in the Methods section, we partition each recognition decision into time bins and train classifiers either on individual time bins or on a cumulatively increasing number of time bins. If brain activity reflects different evidence signals that contribute independent information at different time points, then performance of a classifier trained on neural features from multiple time bins should exceed that of a classifier trained on features from a single time point, since it is able to capitalize on the information from distinct evidence signals (top panel of Figure 1C). If, on the other hand, the neural evidence signal integrates information from all sources, then the signals from previous time points do not contain additional information. Thus, we would expect no benefit for classifiers trained on neural features from multiple time bins in that case (lower panel of Figure 1C).

## Materials and methods

### Participants

The current data set of 132 participants is a subset of the data set for which we previously presented analyses of overt responses (Weidemann & Kahana, 2016; basic analyses of recognition accuracy and response times are repeated here for this subset). Each participant provided informed consent and all procedures were approved by the Institutional Review Board of the University of Pennsylvania. We selected those participants who completed 20 sessions of various free recall tasks. The availability of data from 20 sessions for each participant enabled us to train statistical classifiers on individual participants’ data from 19 sessions (holding out data from one session for cross-validation of classifier performance). As described below, we did this repeatedly such that data from each session was held out once (a leave-one-session-out cross-validation procedure) and all classifier performance measures are based on these held-out sessions only. This yielded enough data to train non-linear classifiers even in cases where not all trials contributed to the classification (as detailed below, some of our analyses placed restrictions on response times).

### Experimental task

As part of a large-scale study of episodic memory, we asked participants at the end of each of 20 sessions to make recognition memory decisions and confidence ratings about words that had been presented earlier in the session for study in various free recall tasks. In each session, participants studied between 12 and 16 lists of 16 words that were each presented for 3 s followed by 0.8–1.2 s of interstimulus interval. In some sessions participants were asked to provide a size or animacy judgment for each word (see Lohnas & Kahana, 2013, for a detailed description of the methods for these sessions) and some sessions included distractor tasks between items and/or between lists. At the end of each list, participants were given 75 s to recall items in any order. A subset of 80 participants were additionally instructed to say aloud any words that were salient during the recall phases following study lists in 6 of the 20 sessions (i.e., an externalized free recall procedure). Some sessions additionally included a final free recall phase after the recall phase for the last list in the sessions. For this final free recall phase, participants were given 5 minutes to recall items from any of the previous lists (for analyses that condition on recall status, we considered recalls from both phases, but recalls during final free recall phases mostly repeated recalls from recall phases immediately following study lists). The recognition test always followed at the end of the session and consisted of 320 probes of which 80%, 75%, 62.5%, or 50% were targets (i.e., words that had been studied in any of the previous lists in the current session) and the rest previously unstudied lure words. The effect of the manipulation varying the proportion of targets vs. lures was small and is not further analyzed here (as detailed below, the training of classifiers took the baserates of the stimulus classes into account to avoid bias).

Throughout the experiment, we obtained high-density EEG recordings, allowing us to investigate brain activity as it unfolds during processing of a memory probe. Each recognition memory trial consisted of the presentation of a probe word, which required a verbal response to the question of whether the given item had been previously studied. We asked participants to substitute “pess” and “po” for “yes” and “no” when answering this question in order to facilitate determination of response times on the basis of the onset of the verbal response (we excluded trials with response times below 300 ms and above 3000 ms from further analyses). Following each binary recognition memory decision, we asked participants to indicate their confidence in the response on a scale from 1 to 5 with 5 indicating the highest level of confidence and 1 indicating low confidence. Most participants indicated confidence ratings verbally; any reference to response times in this manuscript is with respect to the binary recognition decision and not for the confidence ratings. After each classification and confidence rating response, participants pressed the space bar (the recognition response times were determined on the basis of the onset of the verbal recognition decision, however). Following a uniformly randomly jittered interval between 100 and 200 ms after the space-bar press following the confidence rating, participants received visual and auditory feedback on their recognition decision (automatically generated by custom speech recognition software). A 800–1200 ms (uniformly randomly jittered) blank screen separated the offset of the feedback from the presentation of the following recognition probe.

### Data availability

De-identified data and analysis code used in this study may be freely downloaded from the authors’ websites (http://cogsci.info and http://memory.psych.upenn.edu/Electrophysiological_Data).

### EEG data collection and processing

EEG data were recorded with 129 channel Geodesic Sensor Nets using the Netstation acquisition environment (Electrical Geodesics, Inc.). Cz was used as a reference during recording, but all recordings were converted to an average reference offline. For most recording sessions, the EEG aqui-sition software logged periods where the signal changed by more than 250 *μ*V within a short time period as artifactual. Channels where any such period exceeded 10 min. were excluded from the average reference. The multivariate classifiers (described below) weight neural features on the basis of their respective signal to noise ratios and we evaluated their performance by cross-validating on held-out data. Thus we did not attempt to further identify or remove any artifactual activity for our classifier analyses. However, we did exclude twenty-six electrodes that were placed on the face (rather than the scalp) from further analyses.

For event-related potential (ERP) analyses (Figure 2), we excluded channels with low absolute correlation with all other channels and those with extreme levels of raw or log-transformed variance both from the average reference and from the ERPs (we used *z*-scores of ±3 as exclusion thresholds). Additionally we excluded events with significant eye-movement artifacts from the ERP analyses (exclusion criteria were amplitudes in excess of 3 times the inter-quartile range on channels around the eyes any time from 100 ms prior to probe onset until 1000 ms after probe onset).

**Figure 2.**
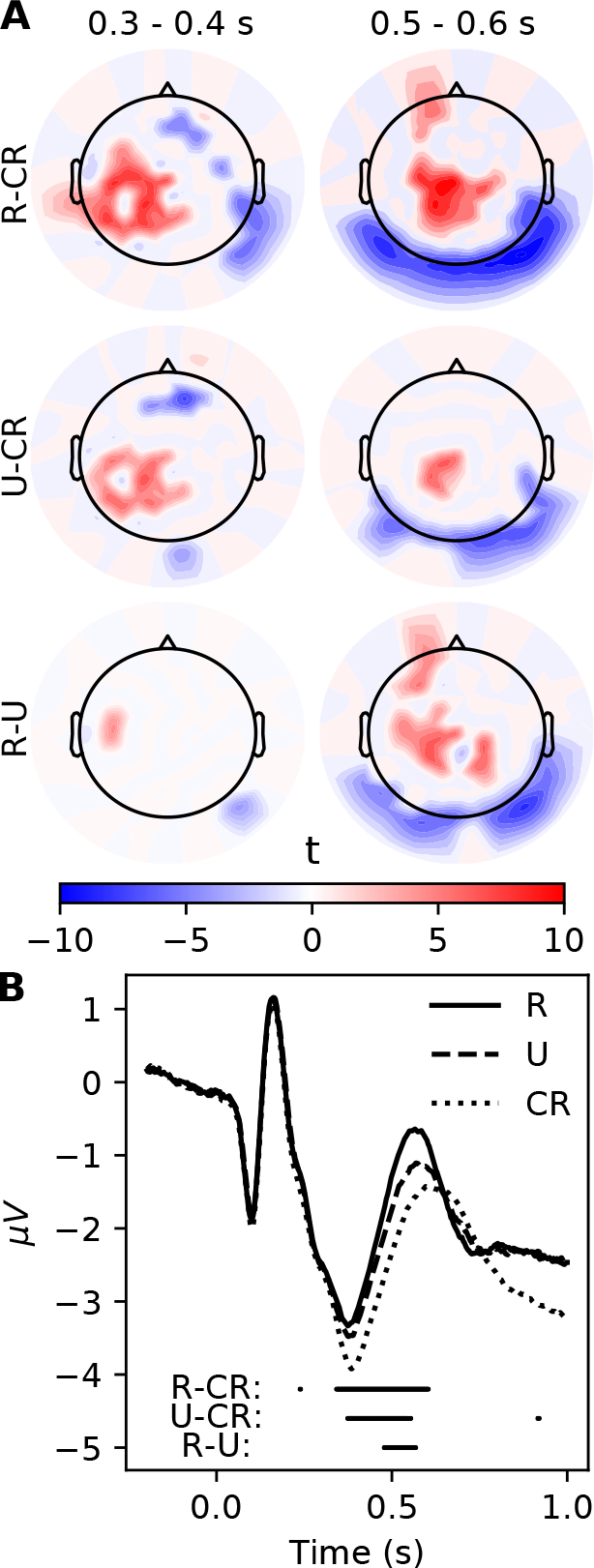
(**A**) Early (0.3–0.4 s) and late (0.5–0.6 s) ERP difference effects for the contrasts between recalled hits (R) and correct rejections (CR; top row), unrecalled hits (U) and CRs (middle row) and between R and U (bottom row). Positive *t*-values (red) correspond to more positive ERPs for R, U, and R trials, respectively. For each contrast we used a permutation test to limit the family-wise error rate across both time bins to 5% and set *t*-values for contrasts that did not exceed the critical *t*-values to 0. For electrodes where all participants contributed to the contrast the corresponding degrees of freedom were 131, but because we excluded data from some electrodes that appeared noisy in a given participant, this number represents an upper bound. (**B**) ERPs for R, U, and CR trials at electrode Cz (on the top of the head). Points below the ERPs indicate time points where the R vs. CR, the U vs. CR, and the R vs. U contrasts were significant which each contrast limiting the family-wise error rate across all electrodes and time points to 5% using a permutation test.

EEG data were partitioned into events starting 500 ms before the onset of a test item and ending 100 ms before the onset of the verbal recognition response. We applied a time-frequency decomposition using Morlet wavelets with 5 cycles for 15 log-spaced frequencies between 2 and 200 Hz, log-transformed the resulting power values, and *z*-transformed these within session. We chose this wide range of frequencies to ensure that we comprehensively capture signals associated with cognitive processes. By separately *z*-transforming power for each frequency within each session, we removed the variability due to baseline power-levels across frequencies and sessions. The multivariate classifier (described in detail below) could then weight power at each frequency, electrode, and time-point according to its predictive value. As is shown in Figure 3, power across all frequencies in the range that covaried with the status of the probe item.

**Figure 3.**
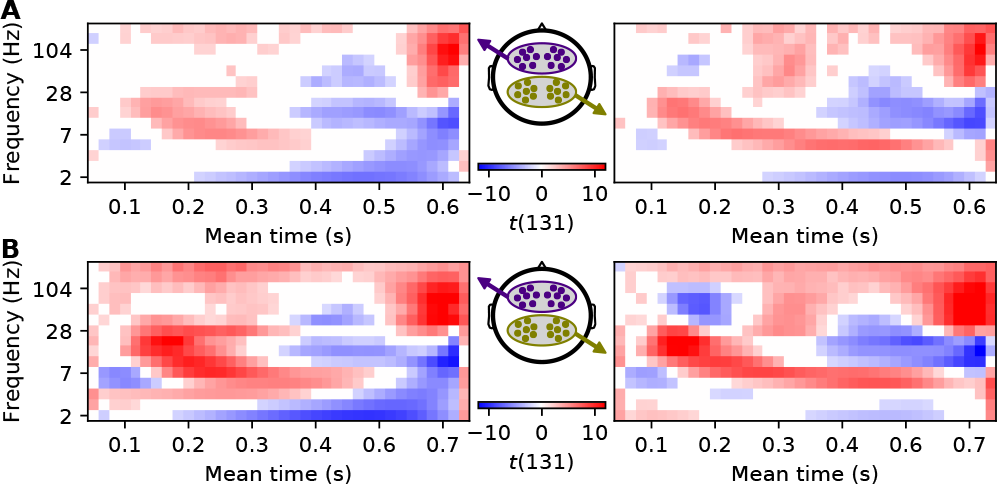
Differences in power for previously recalled vs. not previously recalled hits (**A**) and for all targets vs. lures irrespective of response (**B**). Left and right panels show these differences for anterior and posterior ROIs (illustrated in the middle panels) respectively across frequencies and vincen-tized time bins (mean times associated with some of the time bins are indicated on the abscissas). We used *t*-values for the differences between trial categories for each participant to calculate *t*-values across participants. Red shades indicate higher power for previously recalled hits (**A**) or for all targets (**B**) and blue shades indicate higher power for not previously recalled hits (**A**) or for lures (**B**; within each panel, values that did not reach statistical significance with a false discovery rate of .05 are set to white).

We used a 1500 ms buffer at the beginning of the events and mirrored the last 1500 ms at the end of each event to avoid edge artifacts and to prevent EEG activity from periods during the verbal recognition memory response from bleeding into the analyzed time bins (Cohen, 2014). With the exception of 13 sessions across 5 participants (out of a total of 2640 sessions across all 132 participants) where data were accidentally recorded at 250 Hz, EEG was initially sampled at 500 Hz and down-sampled to 100 Hz after wavelet transformation. We then discarded samples before the onset of the test items, resampled power values for each event to 360 samples, and averaged these samples into 36 equal-time bins for the univariate analyses (Figure 3) and into six equal-time bins for the multivariate classifiers. The lengths of the individual time bins were identical within each trial, but, because response times varied across trials, so did the lengths of the (“vincentized”; Ratcliff, 1979) time bins. We chose to fix the number of time bins to allow us to compare the neural signals across trials as a function of the proportion of each trial’s response time, but we also present some complementary analyses using fixed-length (100 ms) time bins below. To aid with interpretation, whenever reasonable, figures show mean times associated with time bins rather than indicating the corresponding ordinal time bin numbers.

### Classification of EEG data

We used the scikit-learn library (Pedregosa et al., 2011) to train support vector machine classifiers with a radial basis function (RBF) kernel for each participant using a leave-one-session-out cross-validation procedure (all reported classifier results represent the combined performance of 20 models for each participant, where each of these models was trained on 19 lists and tested on the held-out list). The RBF kernel takes the form 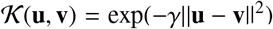, where **u** and **v** are feature vectors and *γ* is a free parameter specifying the width of the Gaussian RBF kernel that determines the region of influence of any support vector. For very large values of *γ* (corresponding to a small variance of the Gaussian kernel) the support vectors are minimally affected by other training examples leading to a highly complex model that is prone to overfitting. Very low values of *γ* (corresponding to a large variance of the Gaussian kernel) make each support vector highly dependent on the entire training set and thus result in highly constrained models. The default setting for *γ* in scikit-learn’s svm.SVC class (using the “gamma=auto” option in its constructor) is to set *γ* to 1/*n_F_*, where *n_F_* is the number of features—we used this setting for all of our fits. Scaling *γ* by the number of features counteracts the increased danger of over-fitting with increasing numbers of features by adjusting each support vectors region of influence to account for larger expected distances between feature vectors as the number of dimensions increase.

The *z*-transformed log-power values across all frequencies and electrodes served as features for the multivariate classifiers. We performed the *z*-transformation separately for each frequency and within each session and thus this transformation was completely separate for training and testing data. These features came either from individual time bins or from varying ranges of time bins starting with the first time bin (bins 1–2, bins 1–3, bins 1–4, etc.). We refer to classifiers trained on features from individual time bins as “individual” classifiers and to classifiers trained on features from multiple consecutive time bins starting with the first time bin as “cumulative” classifiers.

In addtion to the *γ* parameter that specifies the width of the RBF kernel (discussed above), a separate regularization parameter (C in scikit-learn, in other contexts parameters with this function are often labeled *α* or *λ*) determines the width of the margin between the two classes which relates to the tolerance for miss-classifying training exemplars. We used the default setting for this parameter (*C* = 1.0) for all our fits. Because the number of exemplars in each class varied from session to session (as described in the methods the proportion of targets varied between 50% and 80%), we set up the classifiers such that the weights were adjusted inversely proportional to class frequencies (using “balanced” as input to the “class_weight” keyword for the constructor to scikit-learn’s svm.SVC class). This resulted in the regularization parameter (C) for each class, *i*, to be set to *n_S_*/(2*n_i_*), where *n_S_* is the number of samples and *n_i_* is the number of items in class *i*. This prevented the classifiers from being biased towards predicting the most frequenct class.

For the classification of features from 100 ms time bins, we only included trials where responses occurred 750 ms or more after probe onset and only included the 100 (out of 132) participants with at least 30 such trials in each session. Because we considered the time bin starting at probe onset, as well as 11 additional time bins that each had a 50 ms overlap with the previous time bin, this ensured that the last time bin (ending 650 ms after the probe onset) was separated from the response by at least 100 ms. These restrictions did not apply to classifiers using variable-length time bins which we applied to the entire recognition periods until 100 ms before response onset for all 132 participants.

## Results

Traditionally, researchers have averaged voltage time series from EEG recordings to obtain ERPs whose peaks and troughs can be compared across conditions (Luck, 2005). Figure 2A shows “early” (0.3–0.4 s) and “late” ERP (0.5–0.6 s) contrasts for comparisons between correctly recognized targets (“hits”) that were previously recalled (R) and correct rejections (CR), between previously unrecalled (U) hits and CRs, as well as between R and U trials. Even though there is considerable variability in the topography and timing of such ERP contrasts in the literature (Paller, Lucas, & Voss, 2012), the pattern of the contrasts between hits and CRs is broadly similar to that of comparable contrasts in other experiments (Burns, Tree, & Weidemann, 2014; Danker et al., 2008). Consistent with the traditional interpretation of recognition ERPs as reflecting an early familiarity signal and a late recollection signal (Wilding & Ranganath, 2012), ERPs at both time intervals distinguished hits from CRs and the topography of these contrasts differed across these intervals (Figure 2A). Studies of testing effects in recognition memory have suggested that previous recall of an item selectively enhances recollection in a recognition memory test (Chan & McDermott, 2007). To the extent that this conclusion holds, the fact that the difference in ERP contrasts for previously recalled vs. previously unrecalled hits was particularly pronounced for the late interval, further supports the interpretation of late ERP contrasts as reflecting a recollective signal. Figure 2B illustrates this pattern in in individual ERPs for R, U, and CR trials recorded at electrode Cz (at the top of the head): ERPs for hits and CRs separate early and remain separated for hundreds of ms, with ERPs for R and U trials only separating later.

ERPs mainly reflect phase-locked low-frequency power of the underlying EEG activity and are less sensitive to other spectral features that have been shown to reflect cognitive processes involved in episodic memory (Nyhus & Curran, 2010; Jacobs, Hwang, Curran, & Kahana, 2006). For all of our remaining analyses, we therefore decomposed the EEG signal into power across a wide range of frequencies. Additionally later time points in traditional ERP analyses frequently overlap with response periods. To avoid analyzing neural activity associated with executing the recognition response, we only analyzed brain activity up to 100 ms before the onset of this response and used a mirrored buffer that prevented any later brain activity from leaking into the analyzed time period (Cohen, 2014; see Methods for details).

### Power contrasts

Given that previous recall of an item has been linked to selectively enhanced recollection (Chan & McDermott, 2007), we aimed to identify any signals reflecting recollective processes by contrasting spectral power for previously recalled and previously unrecalled hits. Figure 3A shows the dynamic patterns of these contrasts for sensors in two regions of interest (ROIs) that have been frequently used in EEG investigations of familiarity and recollection (Schwikert & Curran, 2014) with both ROIs yielding broadly similar contrasts.

To directly track neural evidence distinguishing old from new items, we also calculated contrasts between spectral power for targets and lures irrespective of the subsequent response. Figure 3B shows that the pattern of these contrasts was remarkably similar to those for contrasts between previously recalled and not previously recalled hits (shown in Figure 3A). Under the assumption that memory is strongest for previously recalled targets, weaker for not previously recalled targets, and weakest/absent for lures, the patterns in Figure 3 could reflect a single memory strength signal that falls out of any contrast between two conditions that vary in memory strength (Squire, Wixted, & Clark, 2007). The fact that these patterns changed quite drastically in the lead-up to the memory decisions, however, might also reflect independent sources of evidence with distinct time courses. An assessment of the relative merits of these alternative accounts, therefore, requires us to quantify the neural evidence in the trial-by-trial variability of EEG activity that distinguishes between targets and lures in the lead-up to a recognition memory decision.

### Quantifying neural evidence

Despite previous efforts to relate the familiarity and recollection components of dual-process models to different (temporally distinct) neural signals (Curran, 1999; Rugg & Curran, 2007), little is known about the actual dynamics of information accumulation in recognition memory decisions and how they relate to accuracy, response times (RTs), and response confidence. Building on the success of machine learning techniques in neural data analyses that have provided unique insights into the dynamics of cognitive processes (Polyn, Natu, Cohen, & Norman, 2005; Norman, Polyn, Detre, & Haxby, 2006; Ratcliff, Philiastides, & Sajda, 2009; Philiastides & Sajda, 2006; Ratcliff et al., 2016; Anderson et al., 2016), we trained statistical classifiers on spectral EEG features to track the neural dynamics of evidence accumulation during recognition memory decisions. A classifier’s ability to distinguish targets from lures can be directly compared to an individual’s recognition memory performance through the use of receiver operating characteristic (ROC) functions relating hits to “false alarms” (incorrect classifications of lures as “old”). The area under an ROC curve (AUC) serves as a convenient index of classification performance, with an AUC of .5 indicating chance performance and an AUC of 1.0 indicating perfect classification (Fawcett, 2006). We previously used confidence ratings and latencies for binary recognition memory decisions to generate ROC functions and showed a strong correspondence between the respective AUCs in the dataset from which the current dataset was derived (Weidemann & Kahana, 2016). Here we assess the evolution of a neural signal indexing evidence for the recognition memory decision by generating ROC functions from the outputs of classifiers that were trained to distinguish targets from lures using spectral EEG features from various intervals during the recognition period. To reduce computational complexity and generate more reliable features for the classifiers, we aggregated the time bins shown in Figure 3 by averaging them in groups of six, partitioning each recognition memory decision into six equal-time bins (see Methods for details).

#### Neural evidence across the entire recognition period

For each participant, we trained a classifier on spectral EEG features from all six time bins to confirm (in held out sessions) that the neural signal in individual trials reliably distinguished between targets and lures (AUC = .71, *t*(131) = 34.59, *SE* = 0.021, *p* < .001; Figure 4A). Single-process models of recognition memory commonly assume that evidence for targets is more variable than that for lures (Wixted, 2007), with converging evidence for this assumption coming from fits of detailed models of evidence accumulation (Starns & Ratcliff, 2014; Starns, 2014; Ratcliff et al., 2016). Larger target variability can result in increased AUCs that are based only on “old” responses (or corresponding classifier output) compared to those reflecting overt responses or classifier output for “new” decisions only (Weidemann & Kahana, 2016), but this pattern is also compatible with dual-process models. Conditional AUCs indicate how much signal the measure of interest contains for each response class (beyond the signal contained in the binary classification of test items as “old” or “new”; Weidemann & Kahana, 2016) and were consistently larger for “old” classifications across all measures (*t*(131) = 8.22–33.22, *SE* = 0.005–0.008, *ps* < .001).

In principle, a classifier trained on neural data to distinguish targets from lures may use different signals than those which are most important for the individual’s recognition memory decision. Indeed, it is unlikely that the coarse measure of scalp EEG activity (compressed into power for a small number of frequencies) could reflect the neural signals leading to the recognition memory decision with high fidelity. In that light, it is of particular interest to what extent the qualitative pattern of (conditional) AUCs is similar across measures. Figure 4B illustrates similarly close relationships between AUCs based on overt responses (AUC_C_ and AUC_L_, with the subscripts denoting confidence ratings and response latency, respectively) and AUCs based on EEG-classifier output (AUC_EEG_; *r* = .71 & .76, *t*(130) = 11.53 & 13.36, *ps* < .001) We also observed strong correlations between conditional AUCs based on overt responses and EEG activity (Figure 4C; *r* = .17–.81, *t*(130) = 1.97–15.58, *p* ≤ .05).

**Figure 4.**
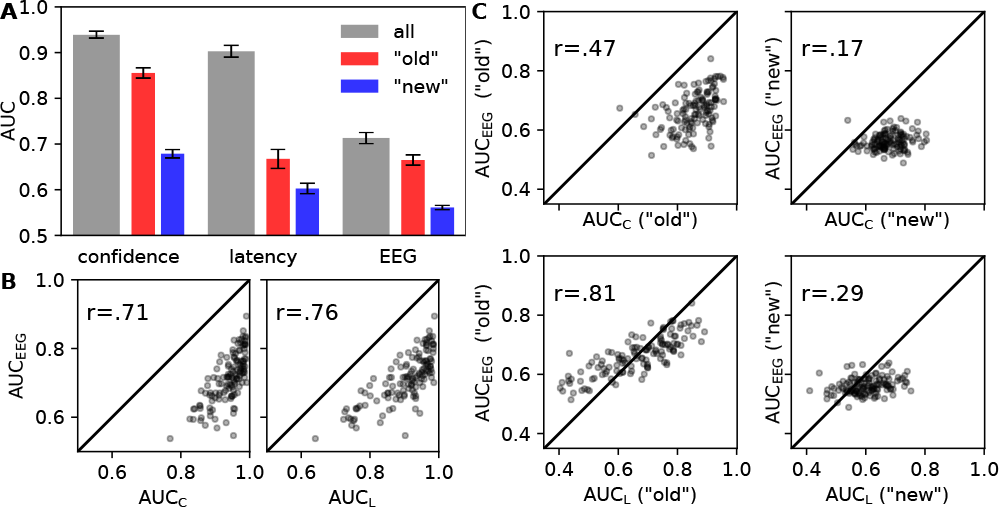
(**A**) Areas under the ROC functions (AUCs) for confidence ratings, response latencies, and EEG activity. AUCs for conventional ROC functions are shown in gray and those for conditional ROC functions based on only “old” or “new” responses (or corresponding classifier output) are shown in red and blue respectively. Error bars correspond to 95% confidence intervals. (**B**) Scatter plots illustrating the relationships between AUCs for either confidence ratings (AUC_C_; left panel) or response latencies (AUC_L_; right panel) and EEG activity (AUC_EEG_). (**C**) Scatter plots illustrating the relationships between conditional AUCs for either confidence ratings (AUC_C_; top panels) or response latencies (AUC_l_; bottom panels) corresponding to “old” (left panels) or “new” (right panels) recognition decisions and EEG activity (AUCEEG) for “old” and “new” classifications. Corresponding correlation coefficients are indicated in the top left of each scatter plot and the main diagonals are shown for convenience.

#### Tracking neural evidence across time

Having established a neural signal that reliably distinguishes between targets and lures and that strongly correlates with recognition memory decisions, we next asked how this evidence accrues over time. If different types of evidence accrue with distinct time courses, this would lead to distinct relative contributions at different points in the decision process. In dual-route models, these signals are assumed to reflect independent sources of evidence and thus should give a distinct advantage to any classifier trained on features from multiple time bins (which would thus be posed to capitalize on both types of evidence to boost performance), relative to classifiers trained on features from a smaller portion of the recognition period. To test this prediction, we compared performance for classifiers trained on features from a cumulatively increasing number of time bins to performance from classifiers trained on features from individual time bins only. Figure 5A shows the corresponding AUCs for these two types of classifiers as a function of time, as well as respective differences (ΔAUC). The AUCs for both types of classifiers were very similar and increased gradually with time (regression slopes and intercepts for both were 0.02 and 0.59, respectively; corresponding *r* values for classifiers trained on individual and cumulative features were 0.52 and 0.53 respectively, *SEs* = 0.001, *ps* < .001). The two types of classifiers are identical for the first time bin and, in the absence of over-fitting, the additional features used by the cumulative classifier for later time bins cannot decrease classification performance relative to the classifiers using features from only one time bin. At the last time bin, the number of features differ by a factor of six for the two classifiers and, even though the features from earlier time bins clearly contained relevant signal (as shown by reliable classification performance for previous time bins), the difference in classification performance between the two classifiers was minuscule (ΔAUC = .003 at the last time bin and < .01 through out). We observed the same pattern of results for fixed-lenght (100 ms) time bins (Figure 5B; regression slopes were 0.006 and intercepts were 0.5 for both classifiers, with *r* = 0.61 for the individual classifier and *r* = 0.60 for the cumulative classifier, *SEs* < 0.001, *ps* < .001; |ΔAUC| < .006). This pattern of results is what would be expected if the classifier output reflected an integrated evidence signal as it accumulates in the lead-up to a decision.

**Figure 5.**
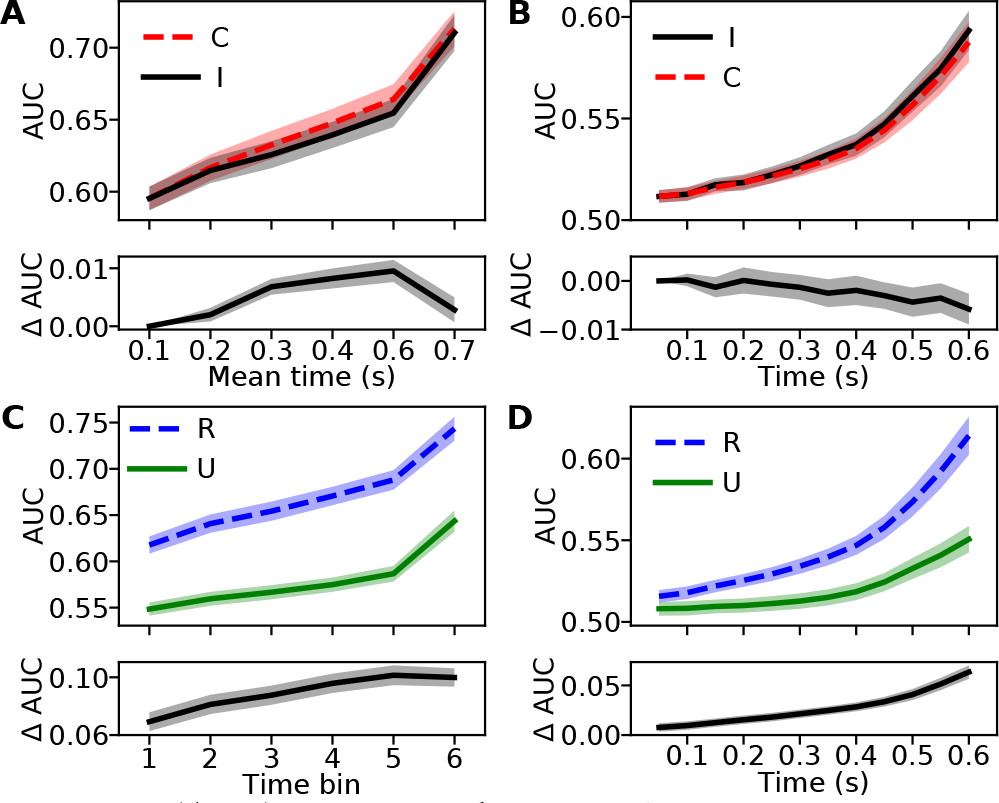
(**A, B**) AUCs for the EEG classifier as a function of vincentized time bin (**A**) or fixed-length (100 ms) time bin (**B**) for features from individual time bins (I) and for cumulatively adding features from each time bin (C; top panels). The bottom panels show the differences between AUCs (ΔAUCs). (**C, D**) AUCs for the EEG classifier as a function of vincentized time bin (**C**) or fixed-length (100 ms) time bin (**D**) when tested on targets that were previously recalled (R) or previously unrecalled (U) and corresponding differences (ΔAUC). Mean times corresponding to vincentized time bins are indicated on the abscissa in (**A**); because RTs differed as a function of previous recall only ordinal time bin labels are indicated in (**C**). Shaded regions correspond to 95% confidence intervals.

To properly assess this evidence against dual-route accounts of recognition memory, it is important to link our neural classifiers to overt responses. Above we compared (conditional) AUCs for memory decisions to those from classifiers trained on neural features from all time bins (Figure 4). Here we track the correspondence between brain activity and overt responses across time by correlating the trial-by-trial output of classifiers trained on individual time bins with subsequent confidence ratings. If classifiers trained on features from different time bins were picking up on different types of relevant signals, we would expect the correlation between classifier output and confidence ratings to peak whenever each signal type provides maximal evidence (e.g., a peak in correlation reflecting an early familiarity signal, followed by another peak reflecting contributions from later recollective processes). From the perspective of a single, continuously accumulating, memory strength signal, however, we would expect gradually increasing correlations as a function of time. Figure 6 shows positive and increasing correlations (measured by Spearman’s *ρ*) for outputs from classifiers and confidence ratings as a function of time (regression slope and intercept were 0.43 and 0.22, respectively, *r* = .56, *SE* = 0.002, *p* < .001).

**Figure 6.**
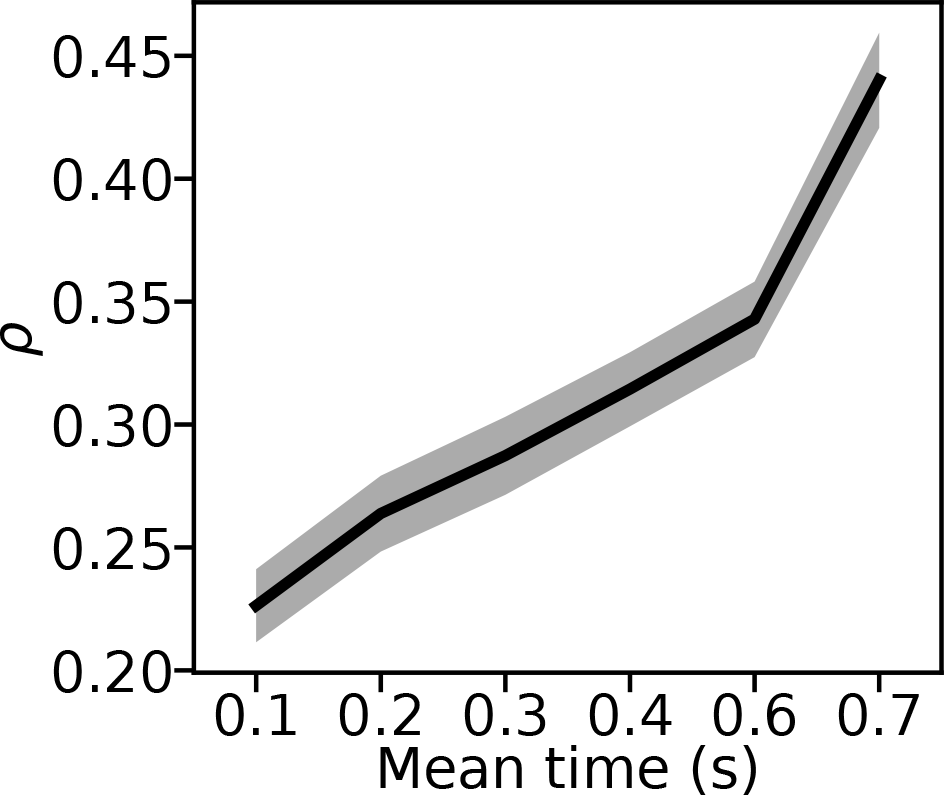
Mean Spearman’s *ρ* between confidence ratings and output of the EEG classifier using features from individual time bins as a function of time bin. Mean times corresponding to vincentized time bins are indicated on the abscissa. The shaded region indicates the 95% confidence intervals.

#### Neural evidence as function of prior recall

Given the converging evidence against dual-route accounts, we set out to maximize our ability to detect any differential contributions from different kinds of processes by assessing the classifiers’ performance conditional on previous recall of targets (Chan & McDermott, 2007). Figure 5C shows that AUCs from recalled trials exceeded those for unrecalled trials across all time bins (*ts*(131) = 21.08–31.19, *SEs* = 0.003, *ps* < .001) and that both types of AUCs increased with time (regression slopes for both were 0.02 with intercepts of 0.54 and 0.61 for AUCs based on unrecalled and recalled targets respectively). Furthermore, the differences between AUCs based on recalled and unrecalled targets also increased with time (regression slope and intercept for ΔAUC were 0.006 and 0.073, respectively, *r* = .27, *SE* < 0.001, *p* < .001). Regardless of previous recall status, classifier performance increased gradually with time as one would expect if evidence accumulated continuously (but at different rates) for both types of trials.

Because differential RTs for targets as a function of previous recall could have contributed to the differences shown in Figure 5C, we also trained classifiers on 100 ms bins of spectral EEG features that we moved from probe onset in steps of 50 ms until the time window that ended at 650 ms after probe onset. Again, AUCs from recalled trials exceeded those for unrecalled trials across all time bins (*ts*(99) = 3.6–19.76, *SEs* = 0.002–0.003, *ps* < .001) and both types of AUCs increased with time (with regression slopes of 0.004 and 0.009 and corresponding intercepts of 0.5 for the AUCs based on unrecalled and recalled targets respectively; corresponding *rs* = .45 & .64, *SEs* < 0.001, and *ps* < .001). Also as above, the differences between AUCs based on recalled and unrecalled targets increased with time (regression slope and intercept for ΔAUC were 0.004 and 0.003, respectively, *r* = .53, *SE* < 0.001, *p* < .001). For both types of analyses, evidence accrual for recalled and unrecalled targets appears most consistent with the continuous accumulation of different amounts (rather than different kinds) of evidence.

## Discussion

Despite a long history of research on recognition memory, there is considerable disagreement about the nature of the evidence that allows us to distinguish repeated encounters from novel experiences (Diana et al., 2006; Dunn, 2004, 2008; Wixted, 2007; Wixted & Mickes, 2010; Squire et al., 2007; Kirwan, Wixted, & Squire, 2010; Dede, Wixted, Hopkins, & Squire, 2013; Yonelinas, 2002; Yonelinas et al., 2010; Malmberg, 2008; Merkow, Burke, & Kahana, 2015; Ratcliff et al., 2016). By training multivariate classifiers to distinguish between previously studied and novel items based on spectral EEG features recorded prior to the execution of a recognition response, we were able to track the accrual of this evidence. We found performance of classifiers trained on a small fraction of the recognition period to be nearly identical to that of classifiers that were able to also capitalize on features from all previous time bins (Figure 5A–B). This suggests that the classifiers directly tracked an evidence signal, rather than a signal over which a decision process integrates to calculate the accumulated evidence—an interpretation also supported by the strong correspondence between classifier output and overt responses (Figures 4 & 6) and by the increasing classifier performance as a function of time (Figure 5). These findings, as well as the strong qualitative similarities between classifier performance as a function of time for previously recalled vs. not previously recalled targets (Figure 5C–D), are difficult to reconcile with dual-route models that posit different kinds of recognition decisions based on independent and temporally distinct familiarity and recollection signals (Reder et al., 2000; Yonelinas, 2002; Diana et al., 2006; Yonelinas et al., 2010).

Our work builds on and extends previous research linking brain activity during an associative recognition task to hypothesized processing stages (Anderson et al., 2016; Borst & Anderson, 2015) and a recent study relating brain activity during a single-item recognition task to parameters in a single-process recognition memory model (Ratcliff et al., 2016). Whereas the former set of studies associated different processing stages with separate familiarity and recollection signals, Ratcliff et al.’s conclusions are largely compatible with ours. Differences in tasks and analysis methods complicate direct comparisons between these studies and our results, but to the extent that the familiarity and recollection signals identified in the partitioning of EEG activity into processing stages (Anderson et al., 2016; Borst & Anderson, 2015) are correlated, they would provide evidence for a unitary recognition signal in our approach which would complement our conclusions (more on this below).

An advantage of our approach is that it does not depend on the assumptions of any specific model of recognition memory, binary choice, or processing stages. By contrasting performance of multivariate classifiers trained on EEG activity during different partitions of the time between probe onset and just prior to the recognition response, we were able to show that “neural evidence” at any given time, incorporates information from previous time points rather than reflecting different kinds of independent evidence signals at different time points. We were also able to directly relate classifier performance and outputs to overt responses, suggesting a close link between neural activity and recognition decisions, as well as associated confidence ratings, that increased as classifiers were trained on later time bins (Figures 4 and 6). As we discuss next, to the extent that we are able to measure all relevant neural signals and to the extent that any signals reflecting independent evidence sources supporting multiple routes to recognition are temporally distinct, our results provide evidence for single-route models of recognition memory.

### Does classifier output reflect recognition evidence?

We have used ouputs of multivariate classifiers trained on spectral EEG activity to track evidence for recognition decisions as it evolved in the lead-up to a response. Our conclusions are conditional upon this approach’s ability to faithfully reflect information that is relevant for the recognition decision. Alternatively, our conclusion that a unitary evidence signal drives recognition decisions could also be due to our approach’s inability to detect neural activity associated with a separate evidence signal. The strong correlations of classifier output with overt responses offer some reassurance by limiting the variance that could be explained by unobserved evidence signals. Additionally, ERP analyses during recognition decisions are often interpreted as reflecting contributions of independent familiarity and recollection signals (Curran, 1999; Rugg & Curran, 2007), which should render these signals observable in our approach.

### Might different evidence signals overlap?

Our analyses depend on different evidence signals exhibiting distinct temporal profiles. Whereas many dual-process models do not specify the dynamics of familiarity and recollection, as discussed above, the near universal assumption is that these signals are temporally distinct. It is generally assumed that an early familiarity signal precedes a later recognition signal, but the fact that responses that are thought to be driven by a recollection signal are made quicker than those thought to rely on familiarity has motivated proposals that recollection occurs before familiarity (Dewhurst, Holmes, Brandt, & Dean, 2006). This pattern of response times, however, is only at odds with an early familiarity and a late recollection signal within the framework of dual-route models. If evidence from all sources is integrated into a unitary evidence signal, as we propose, it is not meaningful to distinguish between “familiarity” and “recollection” responses and we would expect response times to be predominantly a function of overall evidence strength. If memory probes which elicit recollective experiences are generally associated with higher levels of overall evidence (e.g., because recollective evidence is considered particularly diagnostic), single-route models could thus predict faster response times in situations where specific episodic details were recollected, even if recollection signals occur later than familiarity signals.

Our approach is agnostic to the relative order of any familiarity and recollection signals, but to the extent that evidence signals for different routes to recognition overlap in time, we would not be able to distinguish them. Our results are thus compatible with dual-route models, but only under the assumption that recollection and familiarity signals are observable in the EEG activity at similar times or that at least one of these signals does not manifest in the EEG activity. Likewise, if familiarity and recollection signals are highly correlated, we would be unable to distinguish them in our approach. In this case, however, it is unclear to what extent this distinction is meaningful and one could conceptualize these signals as manifestations of a unitary evidence signal instead.

### Novelty detection

Whereas we have considered different ways in which a previously studied item might be recognized as “old”, some evidence suggests that the detection of novelty can also support recognition memory (Daselaar, Fleck, Prince,& Cabeza, 2006; Davelaar, Tian, Weidemann, & Huber, 2011; Kafkas & Montaldi, 2014; Bunzeck, Doeller, Fuentemilla, Dolan, & Duzel, 2009). Our approach quantifies evidence distinguishing between targets and lures and thus is agnostic with respect to whether the relevant signals index familiarity or novelty. It is likely that any familiarity and novelty signals would be strongly (negatively) correlated in standard recognition memory tasks, and some evidence suggests similar temporal profiles for familiarity and novelty signals (Bunzeck et al., 2009). Our approach does not distinguish between strongly correlated and/or temporally overlapping signals and thus is unable to differentiate familiarity and novelty signals with these properties.

## Conclusion

Debates about the relative merits of single- vs. dual-process models often presuppose a false dichotomy between a single route to recognition and contributions from multiple sources of evidence (such as recollection and familiarity) to recognition decisions. Indeed, whereas single-process models assume a unitary “memory strength” signal, models that specify how this value is computed typically derive it from multiple features. For example, the Retrieving Effectively from Memory (REM; Shiffrin & Steyvers, 1997) model represents probes as vectors of features and the recognition decision is based on a computation of the probe’s match to stored memory traces. Thus, the memory strength signal in REM depends on a range of factors (such as the similarity between the probe and stored memory traces, the likelihood of correctly storing features during study, and how features might get confused), but it is considered a single-process model because the effects of all of these factors culminate in a unitary evidence signal. Models that assume separate familiarity and recollection processes, on the other hand, are generally regarded as belonging to a different class of dual-process models, even when they explicitly allow for the associated memory signals to combine into a unitary evidence signal (Rotello et al., 2004; Wixted & Mickes, 2010). As we have argued above, we believe a more meaningful distinction is that between single and dual *routes* to recognition. Specifically this perspective could help resolve apparent conflicts between the success of single process models and evidence suggesting the contributions of distinct sources of evidence to recognition decisions.

There are inherent trade-offs between integrating all available evidence, and thus maximizing the ability to distinguish old from new items, and separately considering different sources of evidence, and thus maximizing the ability to qualify recognition decisions (e.g., with remember-know judgments). It is therefore possible that the extent to which evidence from different sources is integrated into a unitary signal is sensitive to task demands (Rotello et al., 2004; Wixted & Mickes, 2010). At least for the standard old-new discrimination task considered here, however, our results indicate that the memory system integrates available evidence into a unitary evidence signal that drives recognition decisions. These findings thus firmly link recognition decisions to other types of decisions under uncertainty, which are commonly assumed to rely on a unitary evidence signal differentiating between the response options (Ratcliff et al., 2009; Nosofsky, Little, & James, 2012; Philiastides & Sajda, 2006).

